# Assessing the Quality of Cotranscriptional Folding Simulations

**DOI:** 10.1101/2020.01.06.895607

**Authors:** Felix Kühnl, Peter F. Stadler, Sven Findeiß

## Abstract

Structural changes in RNAs are an important contributor to controlling gene expression not only at the post-transcriptional stage but also during transcription. A subclass of riboswitches and RNA thermometers located in the 5’ region of the primary transcript regulates the downstream functional unit – usually an ORF – through premature termination of transcription. Such elements not only occur naturally but they are also attractive devices in synthetic biology. The possibility to design such riboswitches or RNA thermometers is thus of considerable practical interest. Since these functional RNA elements act already during transcription, it is important to model and understand the dynamics of folding and, in particular, the formation of intermediate structures concurrently with transcription. Cotranscriptional folding simulations are therefore an important step to verify the functionality of design constructs before conducting expensive and labour-intensive wet lab experiments. For RNAs, full-fledged molecular dynamics simulations are far beyond practical reach both because of the size of the molecules and the time scales of interest. Even at the simplified level of secondary structures further approximations are necessary. The BarMap approach is based on representing the secondary structure landscape for each individual transcription step by a coarse-grained representation that only retains a small set of low-energy local minima and the energy barriers between them. The folding dynamics between two transcriptional elongation steps is modeled as a Markov process on this representation. Maps between pairs of consecutive coarse-grained landscapes make it possible to follow the folding process as it changes in response to transcription elongation.

In its original implementation, the BarMap software provides a general framework to investigate RNA folding dynamics on temporally changing landscapes. It is, however, difficult to use in particular for specific scenarios such as cotranscriptional folding. To overcome this limitation, we developed the user-friendly BarMap-QA pipeline described in detail in this contribution. It is illustrated here by an elaborate example that emphasizes the careful monitoring of several quality measures. Using an iterative workflow, a reliable and complete kinetics simulation of a synthetic, transcription regulating riboswitch is obtained using minimal computational resources. All programs and scripts used in this contribution are free software and available for download as a source distribution for Linux^®^, or as a platform-independent Docker^®^ image including support for Apple macOS^®^ and Microsoft Windows^®^.

## 1 Introduction

Many aspects of RNA structure and dynamics can be understood quantitatively at the secondary structure level. This includes equilibrium properties of the most stable structure, the existence and equilibrium distribution of alternative structures or structural elements, the (adiabatic) melting and refolding as a response to changes in temperature, the binding to other RNAs, and to a certain extent even the interaction with other types of molecules. These features can be efficiently modelled and computed because (1) the folding energy of an RNA structure is rather accurately approximated by the standard Turner energy model [23] and (2) efficient algorithms exist to compute the partition function of ensembles of RNA structures [15]. Despite this success, however, not all aspects of RNA folding that are important in practice are captured by equilibrium thermodynamics.

RNA folding is an inherently dynamic process and the details of the folding trajectory are at least occasionally biologically relevant. Metastable states, for example, are sometimes the functional ones, as in the example of the Hok/Sok host killing system [8]. In living cells, a nascent RNA starts folding into stable structures while it is transcribed by an RNA polymerase. The structures formed in cotranscriptional folding have decisive functions, e. g. in the case of transcriptional riboswitches, where they decide whether the transcription is terminated or continued to produce a full-length RNA transcript. The structures formed cotranscriptionally are transient and will continue to refold as the transcript is elongated. The mechanisms underlying the biological function of such RNA elements thus can only be understood in terms of dynamics of the folding process.

Riboswitches and RNA thermometers that act on the level of transcription are of practical interest in synthetic biology. They form a class of fast-acting regulators for gene expression that can react to the presence of small molecules via suitable aptamer components or respond to environmental changes in temperature or ion strength that physically affect RNA structure formation. The rational design of such devices, however, requires not only a detailed understanding of the folding process but also computationally efficient methods to model and evaluate the folding dynamics. The size of RNA sequences and the time scales involved preclude 3D molecular dynamics simulations.

Since the standard Turner model assigns an energy to every RNA secondary structure, it is possible to simulate the dynamics of RNA at this level [6]. The calculation and analysis of a large number of trajectories is still computationally expensive and is infeasible at this level when (re)folding processes with time scales of seconds or even hours are to be studied. To be of practical use, e. g. in design applications, furthermore, computational models not only need to provide reasonably reliable predictions, but they also must be much faster and cheaper than experimental approaches. Such methods are indeed available. Instead of simulating the dynamics of RNA secondary structures directly, the idea is to use a coarse-grained representation of the underlying high-dimensional energy landscape and to approximate the (re)folding dynamics of the RNA by a dynamical system on the coarse-grained landscape [27].

A natural coarse graining of an energy landscape is its barrier tree, which only considers the local minima (as representatives of low-energy basins) and the barriers between them [7]. By construction, the barrier tree does not retain any information on the proximity of different basins. Somewhat less coarse abstractions of landscapes, for instance basin hopping graphs [10] or the gradient descent partition of the landscape discussed below, may serve as refine models. On a coarse-grained representation, we can define an (approximate) Markov process that describes the dynamics in terms of transitions between the coarse-grained macrostates. The main advantage of the coarse-grained representation is that the number of macrostates is small enough to explicitly compute the probability of occupancy for a given time instead of having to resort to simulating individual trajectories. As a consequence, the folding dynamics can be computed efficiently for arbitrary time scales – albeit of course only approximately and in terms of the coarse-grained macrostates.

Cotranscriptional folding cannot be modeled by a single energy landscape since the underlying RNA sequence changes with the addition of each nucleotide. The idea of BarMap [9] is to use a sequence of (coarse-grained) landscapes, one for each step of transcriptional elongation. The refolding dynamics between elongation steps is then modeled as a Markov process on the fixed landscape 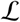. Upon elongation the landscape changes to 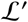, the landscape of the RNA with one additional nucleotide at the 3’-end. Each structure *x* in 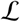, including the local minima of 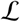, corresponds to a structure *x*′ in 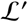 obtained by appending the newly added, unpaired nucleotide. On the other hand, *x*′ belongs to some basin in 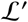 that is represented again by a local minimum. In this way, each local minimum (basin) of 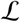 is assigned to a local minimum (basin) in 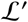, thus defining the “bar map” of Hofacker et al. [9], which relates the barrier trees of consecutive landscapes. In particular, this allows to transfer the probabilities of the coarse-grained macrostates from one landscape to the next. This yields an approximate time-course of the occupancy of coarse grained macrostates for cotranscriptional folding without the need for expensive simulations of individual trajectories. The separation of computations on individual landscapes and the transition between them, furthermore, makes it easy to explore the effects of variations in the speed of transcription on the formation of intermediate structures.

The BarMap software accompanying the publication of the method [9] is a collection of scripts to facilitate the kinetic analysis of several related RNA landscapes, for example under changing environmental parameters such as temperature, or, more relevant here, for a step-wise elongated transcript. It makes use of Barriers [7] to coarse-grain the given RNA landscape, and Treekin [27] to compute the folding dynamics at the level of macrostates represented by gradient basins of secondary structures. The script then stitches together the output of these tools to obtain a result for the full process. In contrast to other simulation tools, BarMap does not rely on the sampling of individual trajectories, but offers an analytical solution based on the enumeration of representative states.

Despite its flexibility and extensibility, the original version of BarMap is a proof-of-concept rather than a ready-to-use tool. Its major drawbacks are:

1. The user has to call multiple scripts to get initial results, making the method unattractive for batch processing even for short input sequences.
2. The output is spread across multiple files, some of which are hardly human-readable, making the interpretation of the results a tedious and error-prone task.
3. There is no feedback that would allow the user to judge the quality of the produced results or help to improve on them.
4. Required software dependencies, in particular Barriers and Treekin, are developed for Linux. Although they can be compiled for other operating systems, this excludes many potential users.
5. There is no comprehensive documentation that guides the user step by step through a typical run, so getting started with the software is unnecessarily complicated.

It therefore takes considerable effort to use BarMap in the context of cotranscriptional folding. Its practical applications there have remained limited.

The intention of this contribution is to alleviate all of the mentioned deficiencies. We give detailed instructions to analyze a real-world example that the reader can easily reproduce on an average desktop computer. Further, we provide a pre-configured working environment including all required software to immediately jump into a cotranscriptional folding analysis. It is deployed inside a highly portable Docker image that can be loaded with a single command and requires no setup apart from a standard Docker installation, which is available on Windows, macOS and Linux. The included scripts can produce preliminary output with just a single command and an input sequence, generating a full cotranscriptional folding simulation and post-processed plots in various formats including PDF and SVG. In addition, quality statistics are computed that allow the user, together with powerful helper scripts, to semi-automatically improve the simulation results if necessary. Finally, an integrating command line viewer for the output files is available that greatly simplifies the interpretation of the results and, together with the generated plots, provides a deep insight into the folding process.

## 2 Theory and programs

A given RNA molecule can, in theory, adopt any of an enormous number of possible structural conformations. It has been shown that, often, sufficiently precise predictions of an RNA’s function can be made by limiting the analysis to its space of *secondary structures*, i. e. structures that are formed by intra-molecular Watson–Crick base pairing according to the following rules: i) each base can pair with at most one other base, ii) base pairs cannot cross each other, i. e. there are no pseudo-knots and iii) paired bases have a minimum distance of three nucleotides. Note that tertiary structure elements, e. g. helix–helix interactions such as in kissing hairpins and other types of pseudo-knots as well as base triples, are not considered. The set of all possible secondary structures *X* for a given RNA sequence s is called the *structure ensemble* of *s*.

### Probability of RNA structures

Each secondary structure *x* ∈ *X* can be assigned an energy Δ*G*(*x*) ∈ (−∞, ∞). Since *x* is an approximation for a *set* of 3D conformations, Δ*G*(*x*) is a Gibbs free energy and consists of both enthalpic and entropic contributions. As a consequence Δ*G*(*x*) is explicitly temperature dependent, a fact that is important, e. g., in the context of modeling RNA thermometers. In practice, Δ*G*(*x*) can be computed in very good approximation from the standard nearest neighbor energy model (Turner energy model), which comprises (temperature-dependent) energy parameters based on detailed thermodynamic measurements [14, 13, 23].

For an equilibrated RNA at fixed temperature *T*, the structure ensemble follows a Boltzmann distribution, i. e. the probability density of *x* is given by its *Boltzmann weight* Z[*x*]: = exp(−Δ*G*(*x*)/(*RT*)), where *R* is the universal gas constant. The *partition function*

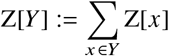

is the sum over all Boltzmann weights for a given subset of structures *Y* ⊆ *X*, and Z: = Z[*X*] refers to the partition function of the full ensemble. The probability of *x* is then given by Pr[*x*] = Z[*x*]/Z, and for a set *Y* of structures, by Pr[*Y*] = Z[*Y*]/Z. The lower the energy of a structure, the higher is its associated Boltzmann weight and thus its probability. We refer to the value Pr[*Y*] as the structural *coverage* of *Y* with respect to the ensemble *X* because it measures how well the subset *Y* covers the most probable (i. e. most important) structures of *X*.

### RNA energy landscapes

For a model of RNA folding kinetics, one needs, in addition to the structure space given by the ensemble *X*, a notion of *neighborhood* to define which structures a given structure *x* ∈ *X* can refold into within an elementary simulation step. Here, the set of neighbor structures *N*(*x*) of *x* is defined as the set of all structures obtained by removing or adding a single base pair from or to *x*. Together with the Gibbs free energy, these ingredients define the RNA energy landscape 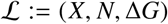.

For any two structures *x*, *y* ∈ *X*, the *transition rate coefficient r*_*x*→*y*_ can be obtained by applying the Metropolis rule [16]:

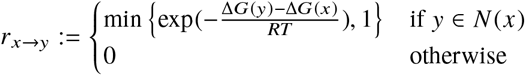

It specifies if and how fast structure *x* refolds into structure *y*. The pair *L*: = (*X*, *r*_.→._), which implicitly determines *N*(*x*) (because *y* ∈ *N*(*x*) if and only if *r*_*x*→*y*_ ≠ 0), represents the *Markov process* induced by the Metropolis rule on 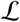. *L* can also be interpreted as a graph where each structure is a node, labelled by its energy, and a weighted, directed edge connects each pair of adjacent structures (*x, y*) such that the edge weight corresponds to the rate coefficient *r*_*x*→*y*_. The weighted graph *L* is connected since each structure can refold into any other one, e. g. by first opening all of its base pairs, and then closing all base pairs of the target structure, one after another.

### Enumeration of the energy landscape: RNAsubopt

The number of secondary structures |*X*| increases exponentially with the sequence length |*s*| of the given RNA [20]. In practice, enumerating the full ensemble is therefore infeasible even for small molecules. A first remedy is to only enumerate those structures with the lowest energies (and, thus, with the highest probability) up to a given threshold Δ_G__enum_, cf. Fig. 1. This can be achieved by applying Wuchty’s algorithm [28] as implemented in the tool RNAsubopt of the ViennaRNA package [12]. The proper choice of Δ*G*_enum_ is non-trivial; it will be used as the main parameter in this work to obtain high-quality simulation results at the lowest possible resource cost. Beside losing possibly important transient structures, the major drawback of this approach is that it may disconnect the landscape, e. g. because all neighbors of a structure lying slightly below the energy threshold are pruned, leaving an isolated state. Please refer to Fig. 1 for an example.

**Fig. 1.**
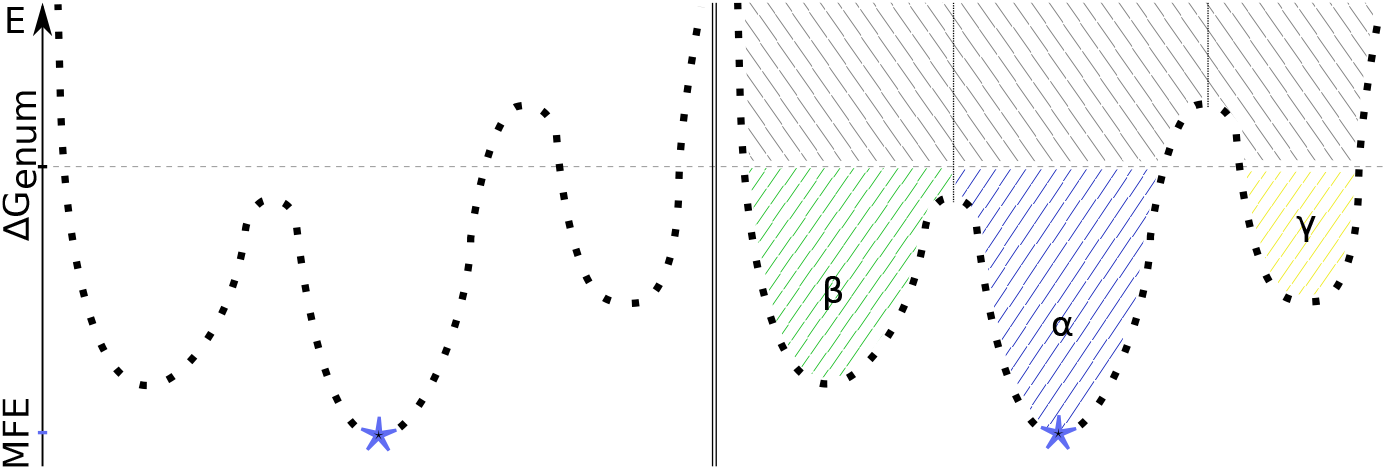
Schematic and strongly simplified visualization of an RNA energy landscape to which a coarse graining is applied. *left:* Individual structures are denoted as black squares, each being adjacent to its left and right neighbor. The *y*-axis displays their free energies. The MFE structure is marked with a blue asterisk. *right:* Coarse-grained version of the same energy landscape. The individual structures have been binned into gradient basins, denoted *α*, *β*, and *γ* here (hatched in green, blue, and yellow, respectively). Due to the enumeration threshold Δ*G*_enum_ (horizontal, dashed line), the upper part of the landscape (hatched in gray) is not part of the coarse-grained representation. As a result, *γ* is disconnected from the remaining basins.

### Coarse graining of the energy landscape: Barriers

Despite the reduction of structures achieved by applying Wuchty’s algorithm, the number of secondary structures is still far too high to use them directly as state set when performing kinetic folding simulations. Therefore, Barriers [7] performs a coarse graining of the structure space *X*. It processes the *sorted* list of low-energy states *X*′: = {*x* ∈ *X* | Δ*G*(*x*) ≤ Δ*G*_enum_} generated by RNAsubopt and extracts (i) the representative set 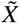 of local minima in *X* and (ii) the energy barriers between them. Barriers also (implicitly) assigns each *x* ∈ *X*’ to the local minimum *γ*(*x*) that it reaches by performing a gradient descent in 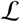, thus binning the structures into well-defined macrostates. Macrostates are therefore simply sets of “microstates”, i. e. individual RNA structures, that form a partition of the landscape. Figure 1 shows the result of applying the described procedure to an example.

For every pair of macrostates *α, β* ⊆ *X*, the rate coefficient for the transition from *α* to *β* is computed as a weighted sum over the rate coefficients of each pair of microstates *x* ∈ *α* and *y* ∈ *β*:

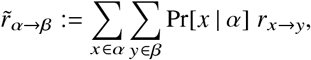

where Pr[*x* | *α*] is the probability that the Markov chain *L* is in state *x* given we know that it is in the macrostate *α* to which *x* belongs. Assuming that basins are steep, the Markov process will approximately equilibrate within *α* before leaving the basin, justifying the approximation Pr[*x* | *α*] = Z[*x*]/Z[*α*]. The approximation fails in particular for large, shallow basins, which are likely to appear in landscapes of sequences with extremely biased nucleotide distributions and unusually large fractions of unpaired bases in their groundstate. The rate constants 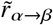 define a Markov chain on the set of basins, which again can be seen as a graph 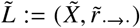 whose neighborhoods are given by 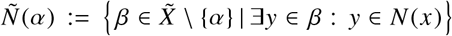. Along with the basins and the corresponding rate matrix, Barriers computes several statistics on the number of structures and the partition functions Z[*α*] for the basins 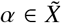.

### Analysis of the folding kinetics: Treekin

Throughout, we consider the folding kinetics of a single RNA molecule as a Markov chain determined by the transition rates *r*_.→._ at the level of individual secondary structures or 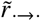 at the level of energy basins as macrostates. Given a distribution *p*(0) of initial states and the rate matrix **R** = (*r_ij_*) composed of the respective rate coefficients *r*_*x_j_*→*x_i_*_ or *r*_*α_j_*→*α_i_*_, the distribution after some time *τ* is simply given by *p*(*τ*) = exp(*τ***R**). It can be computed using standard methods of linear algebra, requiring in essence the diagonalization of **R**. The software Treekin [27] is the implementation used for this work. It can easily process rate matrices of a dimension up to a few thousands. The resulting output gives, for each time step, the population density of each state. Treekin expects a connected state space (*X*, *r*_.→._), a precondition that is easily checked by means of a breadth-first search on (*X*, *r*_.→._). As noted above, the full space of secondary structures is always connected. This is not necessarily true for the output of Barriers, since truncation of the landscape (*X*, *N*, Δ*G*) to low energies may consist of multiple connected components separated by energy barriers above the threshold energy Δ*G*_enum_. The software collection of this contribution contains the script `barriers_keep_connected` that truncates any disconnected state from the rate matrix, thus generating Treekin-compatible input. As an alternative, heuristic methods such as findpath from the ViennaRNA package [12] or RNAEAPath [11] could be used to estimate energy barriers and, thus, approximate transition rates between the components of a disconnected landscape. However, this is beyond the scope of this contribution.

### Cotranscriptional folding: BarMap

Several pre-computed, coarse-grained energy landscapes 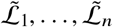 and their associated rate matrices are combined by BarMap to obtain a single kinetics simulation. In our case, 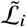 represents the growing RNA sequence of length *i*. To achieve this, BarMap first generates a mapping from 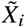 to 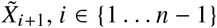, mapping each basin 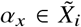, by taking its representative minimum *x* = arg min_*u*∈*α_x_*_ Δ*G*(*u*), appending an unpaired base to obtain *x*′, and then performing a gradient descent [3] on it. This yields *y*: = *γ*(*x*′), a local minimum in *X*_*i*+1_ representing a basin 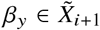.

In practice, the structure *y* does not necessarily represent a basin in 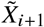, e. g. because the low-energy part 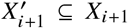 was not enumerated up to sufficiently high energies to recover all previously connected minima, cf. Fig. 2. Barriers also applies heuristics to remove small shallow minima. In both cases, 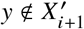 may be the consequence. Furthermore, RNA energy landscapes typically contain adjacent structures with the same energy, thus there is no guarantee that *y* = *γ*(*x*′) is the representative minimum of a basin computed by Barriers. To handle all this, BarMap uses an approximate approach and maps *α_x_* to another 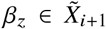 such that the *distance* of minima *y* and *z* is minimal among all macrostates in 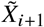. More precisely, the current implementation uses the *base pair distance* defined as |(*y* \ *z*) ∪ (*z* \ *y*)|, i.e. the number of base pairs present in one structure, but not in the other. A basin mapped with a distance of 0 is said to be mapped *exactly*, otherwise it is mapped *approximately*, cf. Fig. 2.

**Fig. 2.**
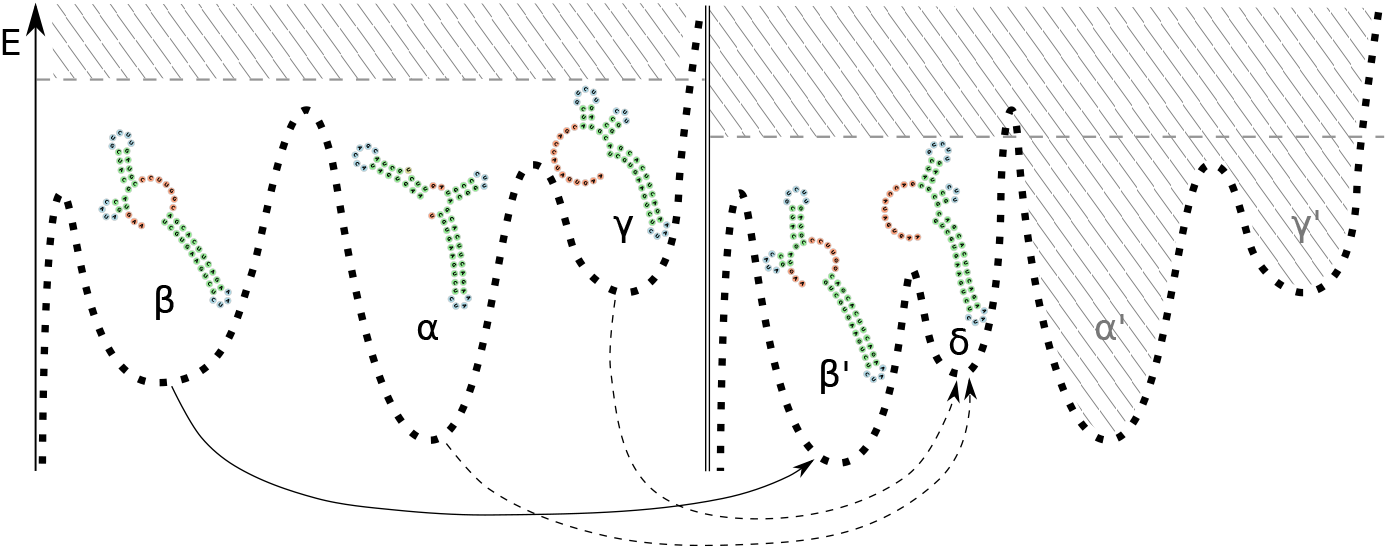
Mapping of some exemplary basins for two coarse-grained, partially enumerated RNA energy landscapes. The left landscape is mapped to the right one. The dotted lines, again, indicate the RNA structures of the individual landscape, and each Greek letter denotes a basin. The structure of each basins’ minimum is plotted above. Solid arrows mark exact mappings while dashed arrows are approximate ones. Basin *β* from the left-hand landscape is mapped to the equivalent basin *β*′ in the right-hand landscape, which has a lower local minimum due to the extension of the 3’ hairpin by one base pair. Since the enumeration threshold (grey, horizontal, dashed line) of the right landscape is lower, the saddle between *α*′ and *β*′ is not enumerated. Thus, *α*′ and *γ*′, the equivalent basins of *α* and *γ*, are now disconnected and removed from the landscape. Instead, *α* and *γ* are mapped to a new basin *δ* due to their low base pair distance to the local minimum of *γ*.

Next, Treekin is applied to the rate matrix of 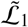 with a defined simulation duration. Then, the final populations of all basins in 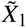 are transferred to the respective basins in 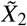 using the mapping defined above. This process is continued until the final landscape 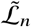 is reached. As a result, a cotranscriptional folding simulation of the RNA molecule is obtained.

### Measuring simulation quality

A major problem of the original BarMap pipeline is that one cannot easily assess the quality of the produced output. To tackle this issue, our software provides rich statistics to evaluate…

1. the *ensemble coverage* of each generated landscape to ensure all highly probable (i. e. important) structures are contained in the simulation,
2. the summed probability density of all exactly mapped (as defined in Paragraph “Cotranscriptional folding: BarMap”) macrostates – *exact coverage* for short – for each mapping step to ensure correctness of the mapping in the case of an equilibrated RNA, and
3. the fraction of *exactly mapped population* during the kinetics simulation to ensure highly populated states are mapped correctly during the simulation.

Here, (1) measures the quality of the individual energy landscapes generated by Barriers, (2) measures the quality of the macrostate mapping generated by BarMap, and (3) measures the quality of the entire kinetics simulation. The coverage of a macrostate *α* is Pr[*α*] as explained in Paragraph “Probability of RNA structures”. The total coverage of all enumerated basis is the ensemble coverage (1), which is directly controlled by the enumeration threshold of Wuchty’s algorithm and serves as a measure of completeness of the coarse-grained landscape. The exact coverage of the mapping (2) is the sum of coverage of all macrostates that are mapped exactly to the next energy landscape by BarMap. Low values indicate that probable structures present in the previous landscape have not been enumerated in the next energy landscape. In such a case, the enumeration threshold needs to be increased for the next landscape and the mapping has to be recomputed. Finally, the fraction of exact population (3) of a mapping step is given by the sum of the populations of exactly mapped macrostates at the end of this simulation step. Here, a low value also shows that an important state was not enumerated in the next energy landscape, which has to be re-enumerated with a higher threshold. The difference is that the inexactly mapped state could as well have a very low coverage, but is highly populated for another reason (e. g. because it is an important transient state, or a kinetic trap).

### Software

The following scripts are provided by our distribution:

- `barmap_gen_barmapfile:` Performs a full cotranscriptional folding run including sequence elongation, energy landscape generation, rate computation, landscape mapping, and output file filtering. Uses a fixed energy threshold for enumerating all landscapes.
- `barmap_rebar:` Used to re-enumerate individual energy landscapes, up to either an absolute energy or by a given energy difference. Also generates transition rate matrices and pre-computes eigenvalues and -vectors of these to speed up the remapping step. Can be run in parallel.
- `barmap_remap:` Recomputes the macrostate mapping and reruns the full kinetics simulation.
- `barmap_filter_treekin:` Used to post-process the final kinetics simulation output. Prunes states below a given population threshold for clearer and more performant plotting.
- `barmap_merge_treekin:` Used to post-process the final kinetics simulation similar to `barmap_filter_treekin`, but additionally merges columns of states that are mapped onto each other in consecutive landscapes. This allows drawing a unichrome curve for a chain of macrostates mapped onto each other despite additional nucleotides being added.
- `barmap_show_run:` Integrated command line viewer for the kinetics simulation output. Used in conjunction with the generated plots to analyze the folding behaviour of the sequence.
- `barriers_coverage:` Computes ensemble coverage and connectedness statistics for the given set of Barriers files.
- `barmap_exact_coverage:` Computes exact coverage values for each mapping step.
- `barmap_exact_population:` Computes exact population fractions for each mapping step.
- `barmap_plot_treekin:` Generates a kinetics plot from the (possibly merged or filtered) Treekin output file using the Grace package [22]. Supports output of PDF documents, SVG vector graphics, and lossless PNG pixel graphics.

The distribution also contains utilities that are not directly important for the example demonstrated here. For a complete list, run `ls /BarMap_QA/bin` inside the Docker container or the corresponding directory in the source distribution.

### System requirements and setup

The software accompanying this work is deployed inside the Docker image `xilef1337/barmap-qa` hosted on Docker Hub. It is downloaded automatically when executing the commands provided in the Method section. The only prerequisite is a working Docker (Community Edition) installation and at least 8 GB of main memory. It supports Microsoft Windows 10 (Pro, Enterprise and Education edtitions), Apple macOS as well as many Linux distributions (e. g. Ubuntu, Debian, Fedora, etc.). Please refer to Docker Docs^1^ on how to install Docker on your platform. Note “Docker installation and configuration”contains instructions on how to avoid common issues encountered during the Docker setup.

Alternatively, a source distribution for Linux is provided on the BarMap-QA homepage^2^. It has been tested on the distributions Fedora 27 and 30 as well as Ubuntu 18.04 LTS. The user is responsible for providing all external dependencies as explained in the `readme` file. Afterwards, the shell script `install.sh` performs all necessary setup steps. We highly recommend to run the test suite by executing the command `make test` inside the distribution directory after the setup script completed successfully. To reproduce the data from this protocol, we suggest using the same versions of all external software, namely ViennaRNA v2.4.13, Barriers v1.6.0, and Treekin v0.4.2.

## 3 Method

In this section, the application of the BarMap-QA pipeline to investigate the synthetic riboswitch RS10 [26] is described. In the original contribution, the authors aimed for designing riboswitches that regulate transcription termination. Therefor, six riboswitch constructs where experimentally investigated and RS10 performed best, showing a 3-fold activation ratio [26]. A cotranscriptional folding analysis of RS10 should show an interesting switching behavior at least between two designed states: (i) the aptamer state, which is able to sense the ligand theophylline, and two (ii) the terminator state, which disrupts the binding-competent aptamer conformation and triggers transcription termination. Following the subsequent step-by-step instructions, the reader will conduct a detailed kinetic analysis of the riboswitch RS10 using default parameter settings.

**Step 1.** We assume that the reader has set up Docker as described in Paragraph “System requirements and setup”. To test your Docker installation, open a terminal (on Windows: a regular PowerShell window) and run:

~~~
docker run hello-world
~~~

You should get a message stating that Docker appears to be running correctly. In case of problems, please refer to Note “Docker installation and configuration”.

**Step 2.** Update and run the `barmap-qa` Docker container. For Linux, we provide the convenience shell script `run_barmap-qa.sh`^3^ that automatically updates the Docker image and starts a new container. Open a terminal running the Bash shell and execute the script. Provide a working directory that is mounted inside the container (here: the current directory “.”).

~~~
./run_barmap-qa.sh.
~~~

For Windows and macOS, the Docker image can be updated by running

~~~
docker pull xilef1337/barmap-qa
~~~

inside a terminal. The container can be started using the following command:

~~~
docker run --rm -v C:\path\to\my\directory:/host -it xilef1337/barmap-qa
~~~

where `C:\path\to\my\directory` is the absolute path to the (existing) working directory of choice. The path to the current directory can be queried using the `pwd` command.

In case the container is started correctly, all generated output will be stored in the mounted working directory of the host machine and will, thus, still be available after the Docker container is shut down. The command prompt should look similar to:

~~~
barmap-qa@489979a0d4ed:host$
~~~

The number displayed in the prompt is a random container id and varies from run to run. The current directory, `host`, should contain all files present in the passed working directory of the host, as can be verified by running `ls`. Finally, create a new sub-directory and change into it:

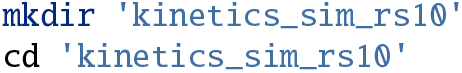

**Step 3.** Now, an initial cotranscriptional folding simulation is performed. Store the RNA sequence in a variable and run the all-in-one pipeline script to generate a complete set of output files:

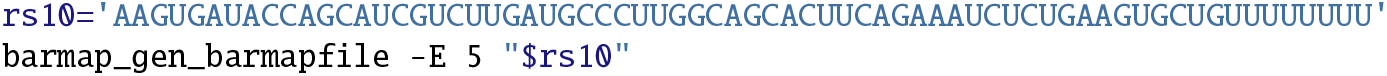

The switch -e sets the enumerated energy range as described in Paragraph “Enu-meration of the energy landscape: RNAsubopt”. There is, however, a technical peculiarity to consider here: -e defines a *relative* threshold ΔΔ*G*_enum_ with respect to the MFE of the current sequence Δ*G*_mfe_ such that the *absolute* threshold is Δ*G*_enum_ = Δ*G*_mfe_ + ΔΔ*G*_enum_.

Here, a threshold of ΔΔ*G*_enum_ = 5 kcal mol^−1^ is used. The lower the value of -e is chosen, the faster the computation and the less resources (computation time and memory) will be consumed. Table 1 in the Appendix lists all generated files and briefly describes their content. The results of the kinetic simulation are automatically plotted and stored in both SVG and PDF format using the batch mode of the plotting tool Grace [22].

**Table 1.** Files that are generated by default when the commands given in method step 3.

| File | Description |
| --- | --- |
| *.bar | Standard output of Barriers containing information about the respective coarse-grained landscape, e. g. representative structure and the barrier height. |
| *.bar.log | Log file that summarizes the Barriers run. |
| *.rates.bin | Binary rate matrix of the respective landscape generated by Barriers. |
| *.evals.bin<br>*.evecs.bin | Eigenvalues and -vectors of the respective rate matrix stored in a binary format. They are used in the final simulation step to speed up the calculation. |
| *.rates.bin.log | Log file of the diagonalization process performed by Treekin. |
| barmap.out | BarMap's state mapping indicating exact (->) and approximate (~>) mappings between consecutive landscapes. |
| barmap.out.kin_t8_1e3 | Full kinetics simulation output generated by multiple consecutive Treekin simulations over all landscapes. |
| barmap.out.kin_t8_1e3.filt | Kinetics simulation output filtered only for highly populated states. |
| barmap.out.kin_t8_1e3.filt.[pdf\|svg] | Plot of the filtered Treekin simulation in SVG and PDF format. |
| barmap.out.kin_t8_1e3.merge | Filtered and merged (cf. above) kinetics simulation output. |
| barmap.out.kin_t8_1e3.merge.[pdf\|svg] | Plot of the merged Treekin simulation in SVG and PDF format. |

**Step 4.** After generating initial results, the quality of the individual energy landscapes can be evaluated. Therefor, it is verified how well an enumeration applying the arbitrarily chosen threshold of 5 kcal mol^−1^ covers the space of all possible RNA secondary structures of RS10. This evaluation is performed automatically by `barmap_gen_barmapfile`, but can also be called manually:

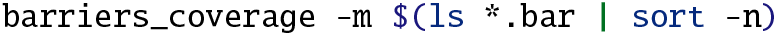

The switch `-m` adds asterisks (*) as markers to entries with low coverage. The notation `$(…)` triggers command substitution and its content is evaluated and inserted. The contained sub-command `ls *.bar | sort -n` lists all Barriers files in the current directory and sorts them numerically such that the growing RNA sequences are processed in the order of their respective lengths. Results are printed on standard out:

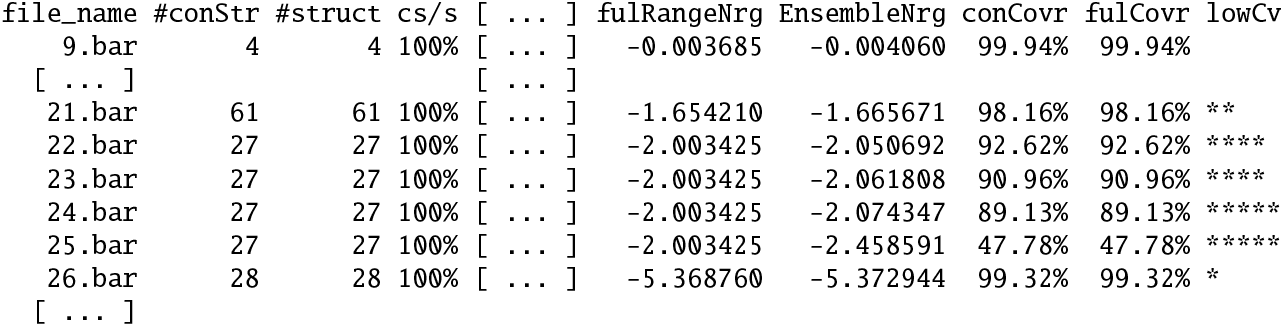

Note that columns or rows omitted due to space limitations are indicated with […] throughout the protocol. The rightmost column, labeled with header the `lowCv`, shows the markers activated by the `-m` option. The more markers are displayed, the lower the coverage is. The exact thresholds at which a certain number of asterisks is given as well as a key to the remaining listed information can be obtained from the help page of the script:

~~~
barriers_coverage --help
~~~

One sees at first glance that many energy landscapes suffer from a bad coverage. Specifically, Barriers files 22–25 (see example output above), 33–37, and 56–60 have less than 95% coverage and are thus marked with at least four asterisks. As an extreme example, file 25 has only 47.78% coverage, i. e. when this sequence folds into a random structure, in the majority of the cases it will not be represented in the generated energy landscape. The potential to observe such a discrepancy increases the longer the sequence gets, cf. Fig. 3.

**Fig. 3.**
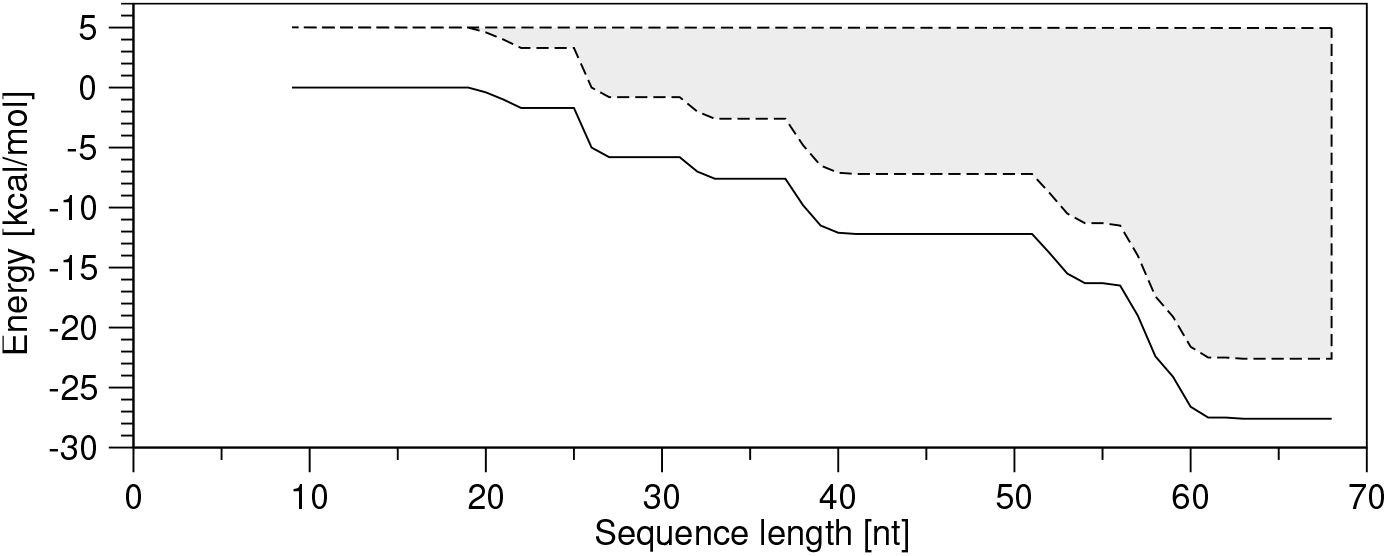
Effect of an arbitrarily chosen ΔΔ*G*_enum_ of 5 kcal mol^−1^ as exploration threshold. The solid black line depicts the energy of the most stable state at each sequence length. As expected, the RNA’s MFE is dropping significantly with increasing length of the transcript. The dashed line above indicates the upper bound of a 5 kcal mol^−1^ energy band and the gray shaded area highlights the increasing amount of potentially missed conformations.

To improve on these results, the exploration threshold ΔΔ*G*_enum_ needs to be increased. For now, it suffices to increase the threshold for *all* sequences by re-running the `barmap_gen_barmapfile` command and changing -e to 8 kcal mol^−1^ (cf. Step 3). Note that by re-running the respective command, all the previously generated output will be overwritten.

Evaluation of the resulting energy landscapes utilizing again the `barriers_coverage` script now shows that an ensemble coverage of at least 97% has been achieved and all landscapes are thus sufficiently well approximated. For an equilibrated RNA molecule, a missing structure will, therefore, not appear with a probability higher than 3% in any intermediate landscape.

**Step 5.** We now proceed to the evaluation of the quality of the “bar map”, i. e. the mapping from each coarse-grained energy landscape to the next one. As before, the quality measures for the mapping are generated automatically by `barmap_gen_barmapfile`, or can be computed manually by executing:

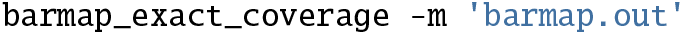

Option `-m` again activates markers for bad mappings, and the file `barmap.out` contains the mappings of all Barriers files. For the previously generated results, enumerating all structures in the energy band of 8 kcal mol^−1^, a dramatic error occurs when mapping Barriers file 25 to file 26. While both individual landscapes expose a high ensemble coverage, a randomly chosen RNA structure will be exactly mapped to the landscape of file 26 in only about 50% of the cases. In the remaining cases, the structure will be mapped approximately to the state of lowest base pair distance, introducing an error into the simulation. In contrast to the original BarMap implementation as described in Paragraph “Cotranscriptional folding: BarMap”, all mappings with a base pair distance less than three are counted as exact mappings in the BarMap-QA pipeline.

The approximately mapped minima are further investigated by re-running the evaluation script and adding the `-a` option to the command. Note that single-letter options can be bundled using a single dash:

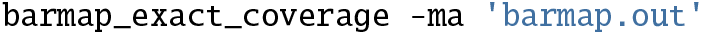

The extended output now summarizes the incorrectly mapped minima together with other useful information like the base pair distance (here 7 base pairs) as well as the saddle height (here −1.70 kcal mol^−1^) with respect to the global MFE basin.

The global MFE basin of file 25, represented by a hairpin structure, is mapped to the open chain conformation in file 26. This mapping completely destroys the characteristic structural feature of the MFE basin and is therefore a bad choice. Additionally, since often the MFE basin has a high Boltzmann weight and thus a high probability, it accounts for 43.89% of the incorrectly mapped probability. This mapping error is fixed by increasing the value of the parameter -e to 9 kcal mol^−1^ when re-running the `barmap_gen_barmapfile` analysis. In the resulting Barriers file 26, the cause of the erroneous mapping becomes obvious (run `head -n6 26.bar`):

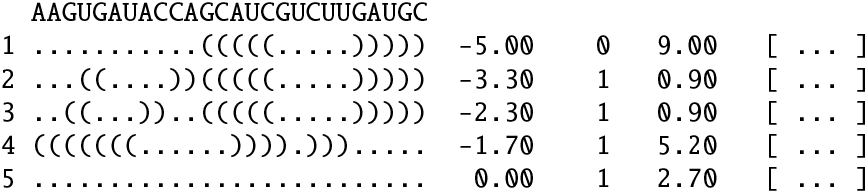

Minimum 4 (−1.7 kcal mol^−1^) is connected to its parent minimum (basin 1) via an energy barrier of 5.2 kcal mol^−1^, such that the saddle structure with an energy of 3.5 kcal mol^−1^ (=−1.7 kcal mol^−1^ + 5.2 kcal mol^−1^) needs to be enumerated to connect this basin to the energy landscape. However, given the global MFE of the sequence from 2 5.bar, −5 kcal mol^−1^, a global exploration threshold of 8 kcal mol^−1^was insufficient as it enumerates only structures up to 3 kcal mol^−1^ (=−5 kcal mol^−1^ + 8 kcal mol^−1^), thus leaving the required structure disconnected in the previous run.

With the increased threshold, the fraction of the exactly mapped probability mass has been increased to more than 99% for most of the mapping steps, with only three steps having 97–98% of exact mappings. Consequently, when using this mapping for randomly chosen structures, there will be a mapping error in less than 3% of the cases.

**Step 6.** In the previous steps, the quality of the mapping has been verified, guaranteeing correctness if each sequence has sufficient time to equilibrate. However, since the whole point of cotranscriptional folding simulations is to observe effects resulting from the population of transient states, this guarantee is insufficient. Therefore, an approach to additionally assess the quality of a simulation run is required.

The script `barmap_gen_barmapfile` also generates population mapping statistics, which indicate how well the populations of the individual states of one energy landscape are mapped to the next one. Again, problematic transitions are marked automatically using asterisks. For the transition from file 26 to 27, one can observe a severe mis-mapping that transfers about 60% of the population incorrectly to the target landscape. For closer examination, the statistics can be recomputed with more detailed information by executing

~~~
barmap_exact_population -ma barmap.out barmap.out.kin_t8_1e3
~~~

where, again, the `-m` switch enables the asterisk markers and `-a` emits every approximately mapped minimum. The issue is quickly identified:

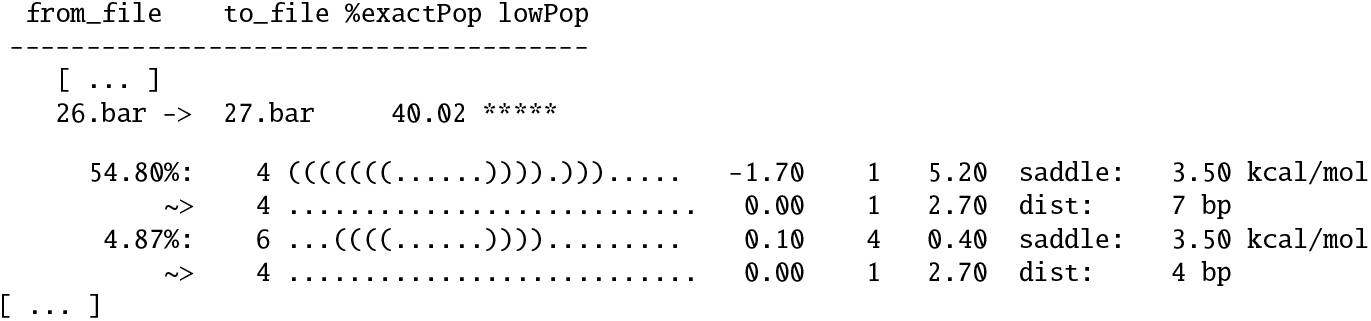

Minima 4 and 6, both exhibiting a leading hairpin structure, are mapped to the open chain. Similar to the problem described above, the exploration threshold ΔΔ*G*_enum_ =9 kcal mol^−1^ is insufficient to connect and explore the saddle point of minima 4 and 6 at 3.5 kcal mol^−1^: in file 27, the MFE is −5.8 kcal mol^−1^, and therefore only structures up to 3.2 kcal mol^−1^ are connected to the landscape with ΔΔ*G*_enum_ equal to 9 kcal mol^−1^. This mapping error did not appear in the last run as the previous version of `26.bar` contained neither basin 4 nor basin 6.

Of course, further increasing the exploration threshold to 10 kcal mol^−1^ and running

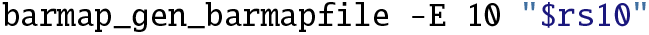

solves this issue for the mapping between file 26 and 27. However, the problem is only postponed to a later transcription step, i. e. the one from file 31 to 32, because the leader hairpin stays populated for a long time period and the MFE value decreases the longer the RNA sequence gets, cf. Fig. 3.

At a first glance, it seems reasonable to simply enumerate all Barriers files such that the saddle point at 3.5 kcal mol^−1^ is included. When looking at the low global MFE of −27.6 kcal for the full length sequence, however, it becomes apparent that this is not a feasible option. A (quite lengthy) run of RNAsubopt enumerating all structures in the the energy range of 31.1 kcal mol^−1^ above the MFE reports a total of about 4.9 billion structures; by far too many to serve as input to Barriers.

### Input size for Barriers

The run time and memory requirements of Barriers strongly depend on the number of input structures. As a rule of thumb, 10 millions of structures may easily be processed, while 50 millions can be processed on a compute server with at least 30 gigabytes of main memory. To count the number of input structures, we refer to Note “Enumeration of structures”. Of course, these values also depend on the sequence length; our example RS10 consists of 68 nucleotides.

Returning to the enumeration of our transient states, we realize the population of these decreased from 54.80% to 19.13% already and will likely vanish within another few simulation steps, so it is sensible to consider a change in strategy. From now on, only some selected energy landscapes will be recomputed while reusing the results for the remaining landscapes. File 32 to file 50 are enumerated up to 3.5 kcal mol^−1^ when running:

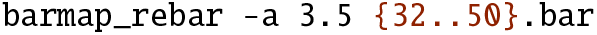

Parameter `-a` is used to specify the *absolute* enumeration threshold Δ*G*_enum_. It is internally converted into the *relative* threshold ΔΔ*G*_enum_ depending on the sequence MFE, which is then used to call RNAsubopt and Barriers. Note that the number of connected minima in file 50 increased from 86 to 2668, thus the execution of the command may take a few minutes. The kinetics simulation is recomputed by running:

~~~
barmap_remap 31.bar
~~~

Here, `barmap_remap` is given the filename of file 31 to start the mapping and simulation process at the corresponding sequence length and to connect the recalculated Barriers files. Since the quality for files 9–30 has already been verified, there is no need to waste resources by repeating these computations over and over again.

As can be verified by another run of `barmap_exact_population`, the quality of the recomputed part of the simulation is already quite good. However, the subsequent mapping step from file 50 to 51 only has an accuracy of about 92%. Most of the incorrectly mapped minima are connected via a saddle at 0.9 kcal mol^−1^. A further accuracy improvement is achieved by enumerating files 51 to 55 up to a higher energy level.

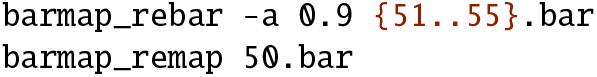

The next drop in exactness occurs when mapping the states of file 59 to file 60.

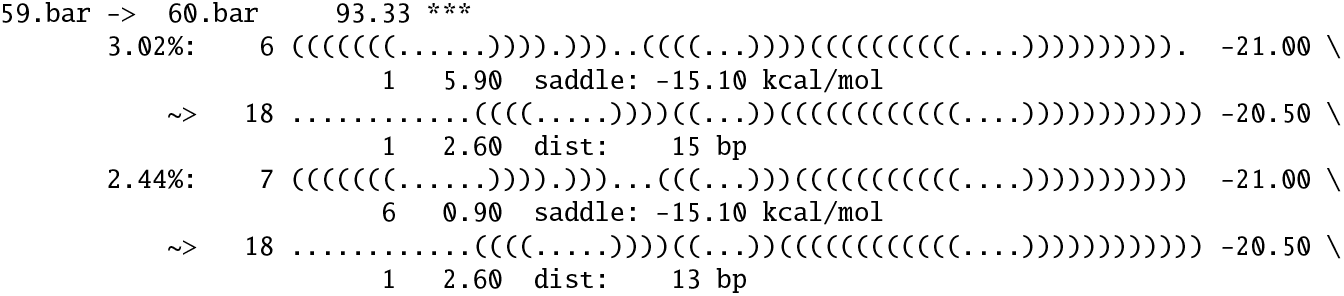

Minima 6 and 7 are connected to the MFE via a saddle structure with an of −15.1 kcal mol^−1^ in file 59. Therefore, file 60 and all subsequent files need to be recomputed. A remapping of the corresponding files to an absolut enumeration threshold of −15.1 kcal mol^−1^ is performed running

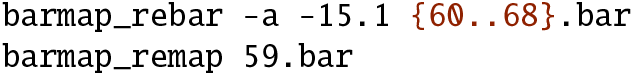

Note that the enumeration threshold of files 56 to 59 is still 10 kcal mol^−1^ and requires no further change, which significantly speeds up the analysis as, e. g., the recalculation of only file 55 already requires 5:24 min on an Intel^®^ Core™ i7-4770 (4 × 3.4 GHz) as well as 1.46 GB of main memory. The sequences in files 60 to 68 share a similar MFE, ranging from −26.6 to −27.6 kcal mol^−1^, see Fig. 3. Here, the change in strategy paid off by making the calculation feasible. A final test of the mapping proofs its exactness. To compute the full kinetics, rerun `barmap_remap` without passing a file name.

**Step 7.** The final results can now be inspected. A line plot is automatically generated using the batch mode of Grace and stored in `barmap.out.kin_t8_1e3.merge.pdf.` An annotated version of this plot is shown in Fig. 4. For similar graphics of all intermediate refinement steps, we refer to Fig. 5 in the Appendix. In such a plot, the *x*-axis shows the time (in arbitrary time units), and the *y*-axis indicates the population density ranging from 0 to 1. Each colored curve is a sequence of basins represented by locally minimal structures that are mapped onto each other by the generated mapping, i. e. the minima of one curve have a similar secondary structure. Additionally, integrated simulation run information can be obtained by applying:

**Fig. 4.**
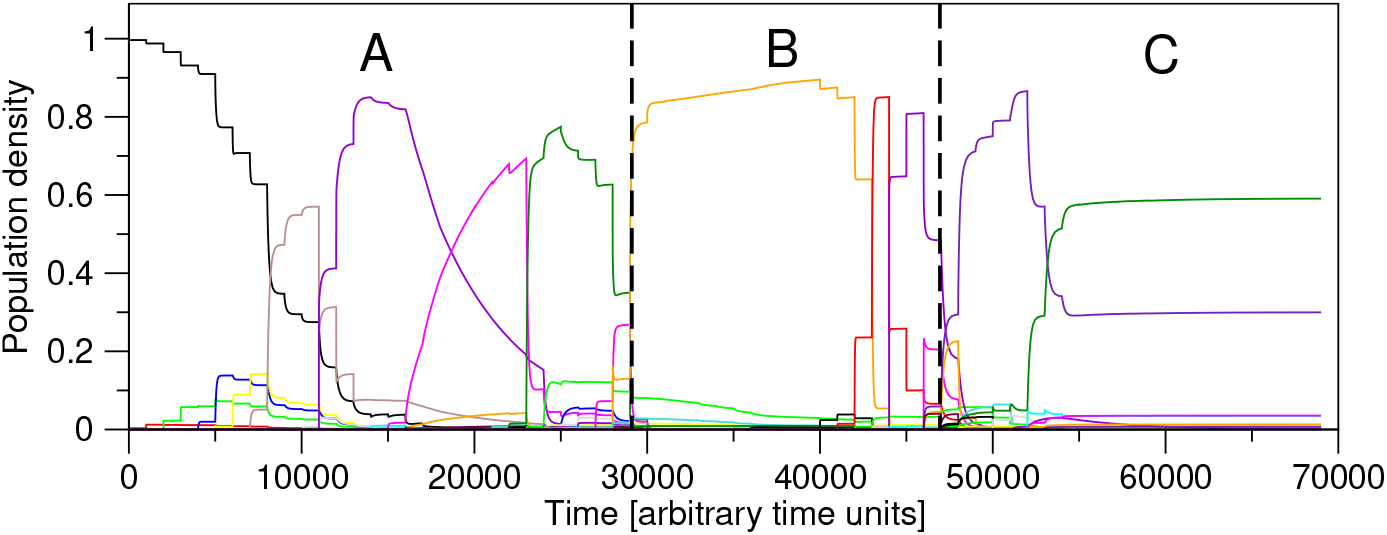
Final results of the cotranscriptional folding simulation of RS10 with BarMap-QA. In conjuntion with the text-based output of `barmap_show_run`, the line plot can be split into three functional parts: *(A)* the transcription and formation of the theophylline aptamer, *(B)* the ligand sensing window, and *(C)* the terminator formation. Although, curves appear and disappear within the latter two these steps are dominated by structures representing the binding-competent aptamer conformation at the 5’-end or the terminator hairpin at the 3’-end of the actual transcript in *(B)* and *(C)*, respectively. Therefore, the actual switching between ligand sensing and terminator formation appears around 47,000 arbitrary time units.

**Fig. 5.**
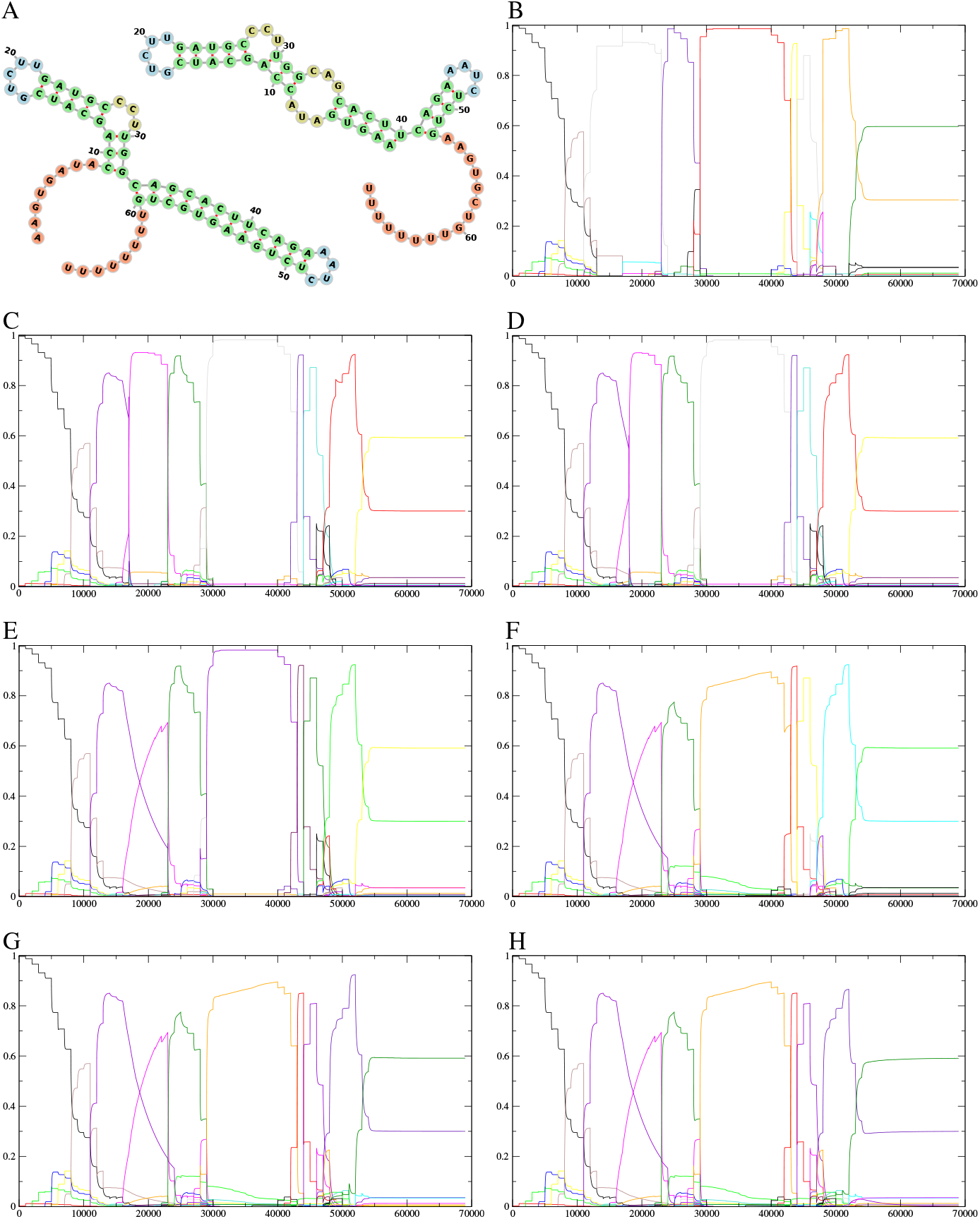
Refinement steps of the folding kinetics analysis of the transcriptional theophylline riboswitch RS10. *(A):* Secondary structure plots of RS10 in its *on* state *(top-right)* and its *off* state *(bottom-left)*. While the *on* state displays the aptamer’s binding pocket consisting of two characteristic interior loops, the *off* state features a stable hairpin loop followed by a poly-U tail, which in conjunction constitute an intrinsic terminator. *(B):* Initial kinetics plot after enumerating all landscapes by 5 kcal mol^−1^. *(C–E):* Kinetics plots after increasing the realtive enumeration threshold to 8, 9, and 10 kcal mol^−1^, respectively. *(F):* Kinetics plot after enumerating Barriers files 32–50 up to 3.5 kcal mol^−1^. *(G):* Kinetics plot after enumerating Barriers files 51–55 upto 0.9 kcal mol^−1^. *(H):* Kinetics plot after enumerating Barriers files 60–68 up to −15.1 kcal mol^−1^.

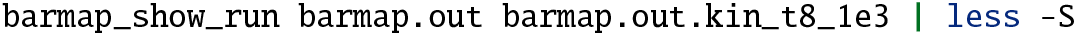

The resulting text-based output summarizes, for each Barriers file, the population densities at the beginning and the end of the corresponding simulation step:

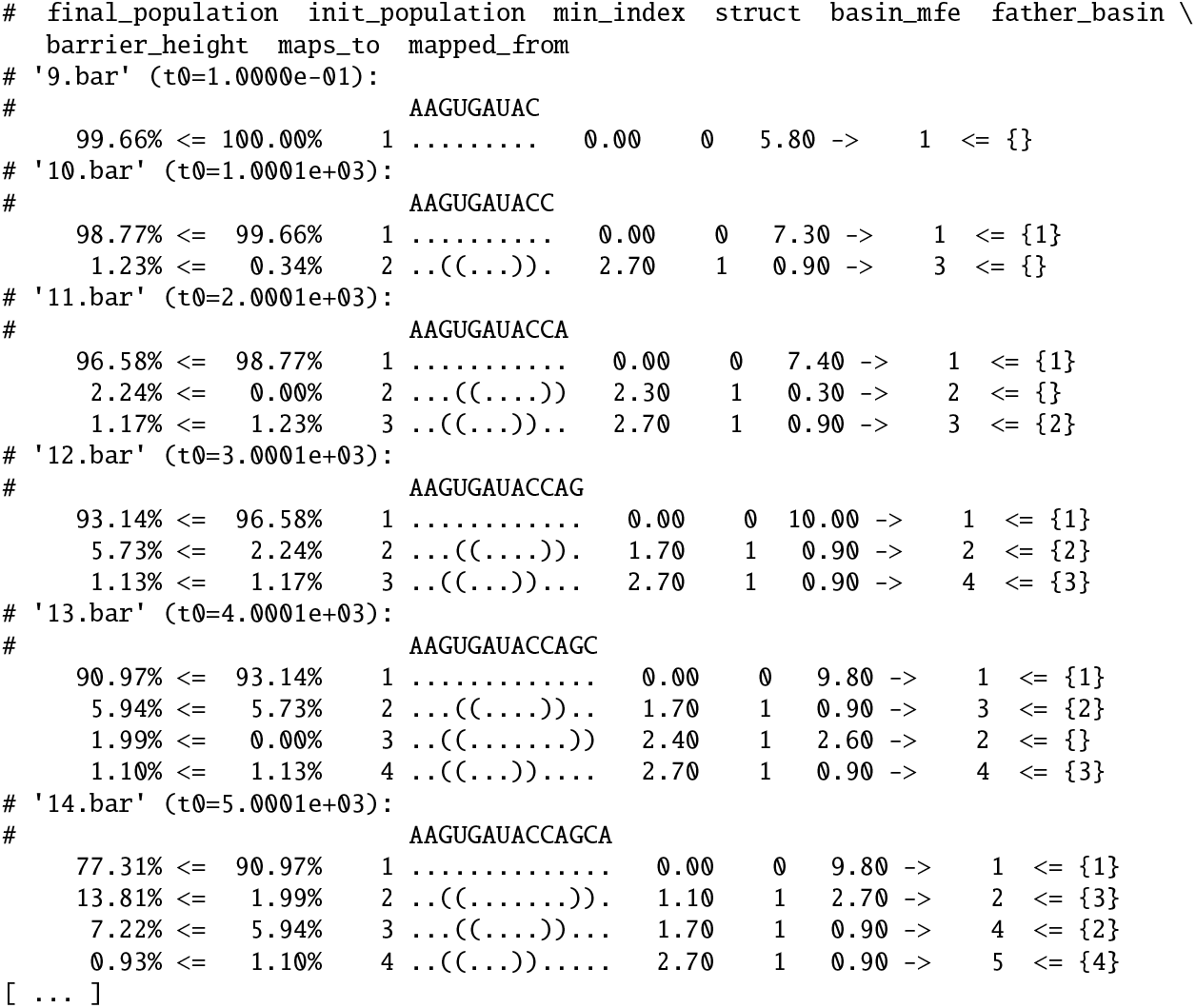

Note that each `t0` indicates the simulation start time in arbitrary time units (au) in the next landscape. Furthermore, for each populated basin the representative structures and the corresponding MFE values are listed. In the example output above, the open chain represents the majority of all populated conformations. In step 14, the corresponding density drops from 90.97% to 77.31% and other, more structured states become populated. In conjunction with the line plot Fig. 4, these information enable the user to easily analyze the generated results.

For the analyzed sequence RS10, the final kinetics simulation shows that as soon as the 42 nt long aptamer sequence is transcribed, the corresponding state is populated with a density of about 86%, cf. Fig. 4 at 33 000 au and Table 2. Although the corresponding curve declines after 42 000 au, other states still containing the binding-competent aptamer conformation at their 5’-end are highly populated. This rapidly changes after 47 000 au when a sufficiently large part of the terminator hairpin is transcribed, cf. Table 3. From now on, the terminator structure gets extended step by step and, thereby, the aptamer structure is suppressed. The exact timing and the speed of switching between the aptamer and the terminator state seems to be essential for functional transcription-regulating riboswitches [26, 25]. A close examination of the supposed switch of states after 51000 au reveals that all the highly populated states contain a complete terminator hairpin and only the structural leftovers of the aptamer conformation differ, cf. Table 3. In other words, when asking for the fraction of states containing the terminator structure, the mass of all the highly populated states could be added up. However, this type of coarse graining is beyond the scope of this protocol.

**Table 2.**
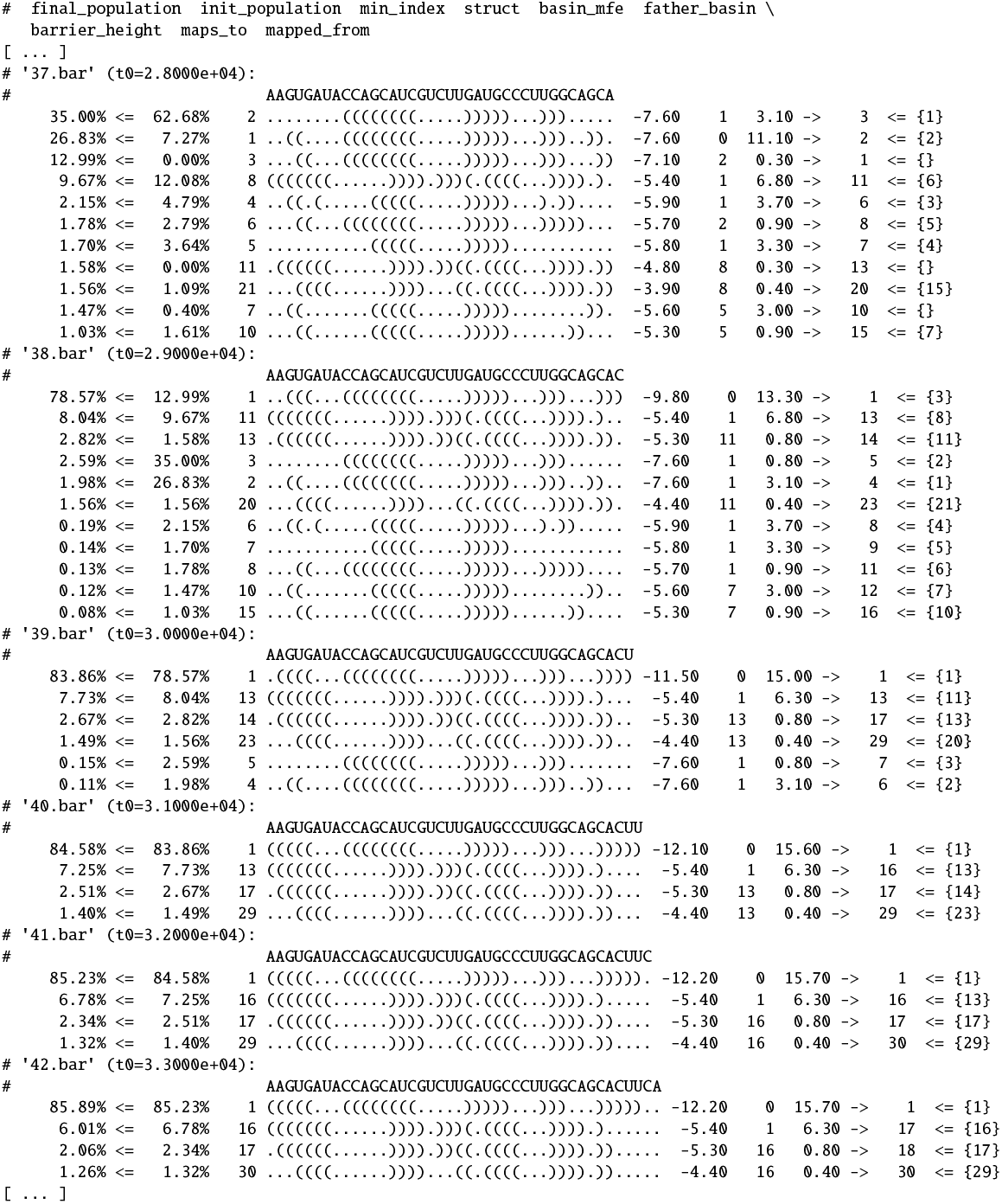
Text-based output for the RS10 riboswitch showing the formation of the binding-competent aptamer structure.

**Table 3.**
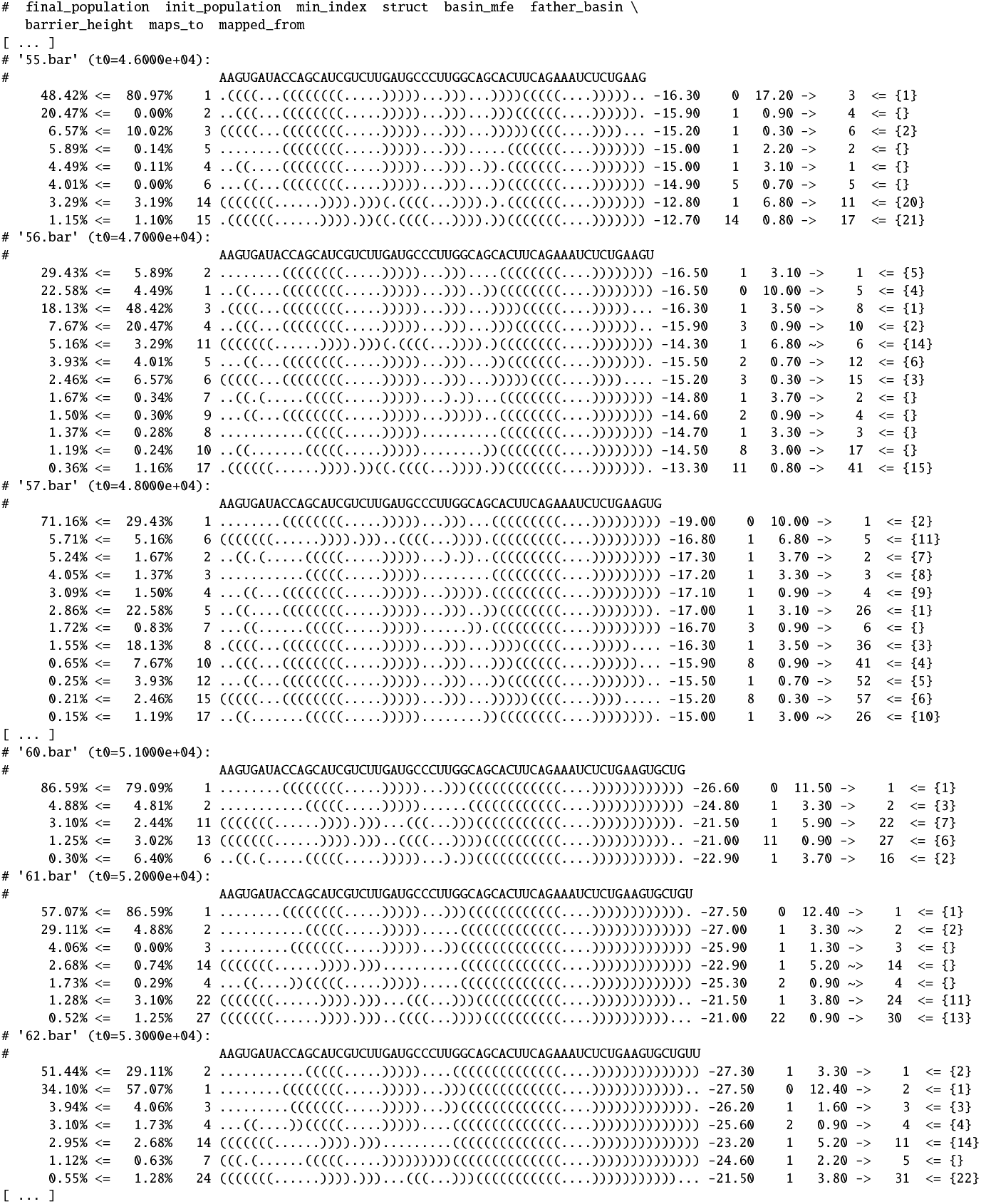
Text-based output for the RS10 riboswitch showing the formation of the terminator hairpin.

## 4 Notes

### Docker installation and configuration

For Windows and macOS, download the Docker software from the corresponding official website. After the installation, you may need to reboot your machine. Ensure the Docker application is started before attempting to use the `docker run` command. On Windows, non-privileged users have to be added to the `docker-users` group to be able to start any Docker containers. On Linux, Docker can usually be installed via the system package manager (package `docker` on Fedora and Ubuntu). Please note that you may have to add a `docker` group (`sudo groupadd docker`) and add the current user to it (`sudo usermod -aG docker “$USER”`) if you get an error when using docker run. Additionally, it may be necessary to configure the system to automatically run the Docker daemon when the system is booting (`sudo systemctl enable docker`). On both Windows and Linux, log out and log in again after adding users to a group to apply the updated settings (or reboot the machine to be on the safe side). If you get “out of memory” or “segmentation fault” errors, add the option `-m 8G` to all docker run commands; it ensures that at least 8 GB of main memory are available to the container. Increase the memory limit if you apply the pipeline to longer input sequences.

### Enumeration of structures

As described in Paragraph “Enumeration of the energylandscape: RNAsubopt”, Wuchty’s algorithm can be applied to enumerate all structures up to a given energy threshold Δ*G*_enum_. However, the exponential dependence of the number of secondary structures on the sequence length imposes practical limitations upon the choice of this parameter, especially if the output is supposed to be processed by computationally demanding methods such as Barriers. Here, we show how to determine the number of structures for a given value of Δ*G*_enum_ such that the user can make a qualified decision about the feasibility of continuing the analysis.

As an example, we consider again the riboswitch RS10, for which a saddle state at Δ*G*_enum_ = 3.5 kcal mol^−1^ had been identified. To connect the respective minimum to the global MFE basin located at −27.6 kcal mol^−1^, ΔΔ*G*_enum_ needs to be set to 31.1 kcal mol^−1^. Running

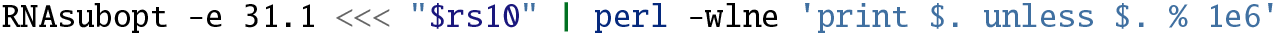

reports about 4.9 billion structures. In detail, the one-line Perl script counts the number of enumerated conformations, indicating its progress every one million structures. %is the modulo operator and `$`. stores the current line number. The switches passed to the Perl interpreter enable warnings (`-w`), automatically add newline characters to printed lines (`-l`), enable line-wise processing of the standard input stream (`-n`), and specify that the script to be executed follows directly on the command line (`-e`).

### Parallelization

The script `barmap_rebar` supports multi-threading. The number of threads to launch can be specified using the switch `-t`. This switch is also supported by `barmap_gen_barmapfile`, which passes it to all other scripts it is calling:

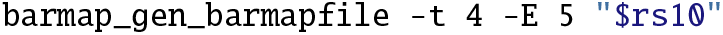

Note that running multiple instances of Barriers with a high number of sequences requires a lot of main memory. We recommend keeping an eye on the spawned threads using a system monitor and setting a per-process memory limit using the shell builtin `ulimit`. For example,

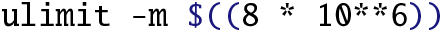

sets the maximum resident set size (i. e. the amount of occupied main memory) for each sub-process of the current shell to 8 GB. The notation `$((…))` triggers arithmetic expansion of its content and ** is the exponentiation operator. The value to `-m` is assumed to be given in kilobytes.

### The influence of transcription rates

Obviously, how close the distribution of structures of a given RNA sequence is to the equilibrium distribution depends on the time available for equilibration per step before the sequence is elongated again, i. e. the rate of transcription. With high transcription rates (i. e. many added nucleotides (nt) per time unit), a transient state may still be highly populated when the RNA chain is extended, and thus a different part of the energy landscape – which could hardly be reached from the ground state – may become accessible. As a consequence, the quality analysis of the kinetics simulation needs to be repeated if the latter is performed with a different transcription rate.

The BarMap pipeline has no option to directly set the transcription rate but, instead, the end time (and thus the duration) of the simulation for each elongation step can be defined in *arbitrary time units (au)*. The question how exactly they correspond to real time is left to the user. Recently, 1 au nt^−1^ has been mapped to 5 × 10^−6^ s nt^−1^ [19], an estimate also applicable to BarMap [1]. Thereby, a transcription rate of 200nt s^−1^translates into 1000 au s^−1^, which is equivalent to 0.005 s nt^−1^. These are only rough estimates, and the true transcription rate varies for different types of RNA molecules [2], environmental conditions such as temperature [18] or growth phase [24]. We therefore advice the reader to tune this parameter to known experimental data, ideally from the biological system of interest.

To change the time per elongation step, the `-e` option of `barmap_remap` can be set:

~~~
barmap_remap -e 4e3
~~~

This changes the default duration of 1 × 10^3^ au to 4 × 10^3^ au, which corresponds to a transcription rate of 50nts^−1^ following the approximations above. Additionally, BarMap-QA adjusts the simulation time of the last landscape to be ten times as long as the the simulation time of all other landscapes. This allows the user to get an impression of the equilibrated structure distribution after the transcription process has finished. Note that, when running `barmap_remap`, only the final BarMap simulation step is repeated. Therefore, previous results are kept and new files with the prefix `barmap.out.kin_t8_4e3` are created.

### Post-processing of the kinetics file

BarMap-QA provides two types of plots, marked with either the `.filt` or the `.merged` suffix, which result from two different post-processing procedures. The first one is a simple filter that removes any state that does not acquire a minimal population from its output, and also allows to limit the total number of states in each simulation step. Manually calling the script `barmap_filter_treekin` allows the user to adjust the minimum population threshold using the `-p` switch. The filtered output can then be plotted using `barmap_plot_treekin`, which supports multiple output formats (currently PDF, SVG, and PNG). Strikingly, the color of the population curves change every time a new Barriers file is processed (i. e. if the next nucleotide is added to the RNA sequence), which can be helpful during the analysis of the plot. The second option is provided by the script `barmap_merge_treekin`. It offers the same filtering options, but additionally merges the states of successive Barriers simulations in the output according to the generated BarMap file. As a result, the population curves have a single color, which often provides a more pleasant visual experience. The output can, again, be processed with `barmap_plot_treekin`.

### Extending BarMap-QA

BarMap is designed to be applicable to a broad range of problems and thus very flexible. Due to the semi-automatic architecture of our pipeline, this flexibility is transparently propagated despite the added functionality. For instance, the user can provide custom rate matrices accounting for ligand binding or changing environmental temperatures, and may still compute the quality statistics as described above.

The current BarMap implementation does not support variations in the rate of transcription along the input RNA – an effect that has been reported in literature [17, 4]. Such a functionality would be desirable e. g. to understand the effect of pause sites [8, 21] on cotranscriptional RNA structure formation. We expect that this functionality will become available in a future release of BarMap.

pourRNA, a memory efficient and parallelizable alternative to Barriers, has been released recently [5]. Integrating this new tool into BarMap-QA might speed up the computation, but remains as a future task.

### Feasibility of cotranscriptional folding simulations

Apart from the length of the input sequence, the feasibility of a BarMap analysis also depends on the sequence’s folding characteristics. An RNA molecule with a smooth energy landscape with a dominant global minimum will be much easier to simulate than bistable sequences exhibiting highly probable alternative conformations or “folding traps”. The reason is that persistent intermediate folding states require an enumeration up to a constant energy level during the entire elongation process, leading to a combinatorial explosion of the number of secondary structures. In contrast, if the most likely folding paths quickly lead to the global minimum, only a small number of structures need to be enumerated to obtain high quality simulation results even for long RNA sequences.

As an example for a sequence that is more difficult to analyze, we encourage the reader to experiment with the non-functional riboswitch candidate RS8 [26], whose sequence^4^ is very similar to RS10.

## Acknowledgements

This work was supported in part by the German Research Foundation (DFG; grants STA 850/15-2 and MO 634/9-2).

We thank Ivo L. Hofacker for fruitful discussions on variations of the BarMap approach, and Anne Hoffmann and Jens Steuck for their IT support to test our software.

## Trademark notices

Windows^®^ is a registered trademark of Microsoft Corporation in the U. S. and other countries. This work is an independent publication and is neither affiliated with, nor authorized, sponsored, or approved by, Microsoft Corporation or other companies. macOS^®^ is a trademark of Apple Inc., registered in the U. S. and other countries. Linux^®^ is the registered trademark of Linus Torvalds in the U. S. and other countries. Docker is a registered trademark of Docker, Inc. in the United States and/or other countries. Docker, Inc. and other parties may also have trademark rights in other terms used herein. Intel^®^ Core™ is a trademark of Intel Corporation or its subsidiaries in the U. S. and/or other countries. All other trademarks are the property of their respective owners.

## List of abbreviations

This work makes use of the following abbreviations:

RNA: RiboNucleic Acid
nt: nucleotide
MFE: minimum free energy
*E. coli*: Escherichia coli
au: arbitrary time units

## Appendix

### Generated output files

When using the all-in-one script `barmap_gen_barmapfile` to automatically run a complete BarMap-QA analysis, all the files listed in Table 1 are generated.

### Visualization of simulation refinements

Throughout the step-by-step instructions described in the Section Method, the kinetics simulation of the riboswitch RS10 is gradually improved. Figure 5 shows plots of all complete kinetics simulations at each step of the refinement process.

### Detailed output for RS10 run

The text-based output of `barmap_show_run` summarizes, for each Barriers file, the population densities at the beginning and the end of the corresponding simulation step. Note that `t0` indicates the simulation start time of each step in au. For each populated basin, the corresponding representative structure and their MFE value is indicated. For a detailed description run `barmap_show_run -h`.

## Footnotes

1 https://docs.docker.com/install/

2 https://www.bioinf.uni-leipzig.de/Software/BarMap_QA/

3 https://www.bioinf.uni-leipzig.de/Software/BarMap_QA/run_barmap-qa.sh

4 AAGUGAUACCAGCAUCGUCUUGAUGCCCUUGGCAGCACUUCACUCCUAGUGGAGUGAAGUGCUGUUUUUUUU

